# Single-cell RNA-sequencing analysis reveals the molecular mechanism of subchondral bone cell heterogeneity in the development of osteoarthritis

**DOI:** 10.1101/2022.03.20.485020

**Authors:** Yan Hu, Jin Cui, Han Liu, Sicheng Wang, Qirong Zhou, Hao Zhang, Jiawei Guo, Liehu Cao, Xiao Chen, Ke Xu, Jiacan Su

## Abstract

The cellular composition and underlying spatiotemporal transformation processes of subchondral bone in osteoarthritis (OA) remain unknown. Herein, various cell subsets from tibial plateau of OA patients are identified, and the mechanism of subchondral microstructure alteration is elaborated using single-cell RNA sequencing technique. We identified two novel endothelial cell (EC) populations characterized by either exosome synthesis and inflammation response, or vascular function and angiogenesis. Three osteoblast (OB) subtypes are introduced, separately related to vascularization, matrix manufacturing and matrix mineralization. The distinct roles and functions of these novel phenotypes in OA development are further discussed, as well as interaction network between these subpopulations. The variation tendency of each population is testified in a DMM mouse model. The identification of cell types demonstrates a novel taxonomy and mechanism for ECs and OBs inside subchondral bone area, provides new insights into the physiological and pathological behaviors of subchondral bone in OA pathogenesis.

## Introduction

Osteoarthritis (OA) is an insidiously progressive, high-cost, and poorly prognostic joint disease, affecting approximately 250 million patients worldwide.^1^ The most common clinical manifestations of OA are chronic cartilage degeneration, subchondral bone microstructure alteration, osteophyte formation, and intractable joint pain.^2–4^ Fundamental research targeting articular cartilage destruction or cell senescence revealed considerable therapeutic potential of cartilage repair methods, however, the clinical trials have failed to varying degrees in recent decades.^5, 6^ Accumulating evidence suggests that pathological alterations inside the subchondral bone are responsible for chondrocyte reduction and matrix degradation.^7, 8^ The turnover rate of subchondral bone remains relatively low under normal circumstances and is accelerated by multiple factors in OA status, including mechanical and biological factors. The uncontrolled bone remodelling in the subchondral bone results in subsequent changes, including hypervascularisation, hyperpathia, abnormal mechanical support, and cartilage destruction.^9^ Currently, both physiological and pathological behaviours of the subchondral bone have been valued in advanced research targeting OA therapies. However, the composition of subchondral bone cell types in patients with OA and the underlying spatiotemporal transformation processes remain unknown.

Mesenchymal stromal cells, osteoblasts (OBs), osteoclasts, endothelial cells (ECs), and immune cells are delicately orchestrated by various biological and mechanical factors in the local microenvironment of the subchondral bone.^9^ Under abnormal loading conditions, the activated form of TGF-β is released from the bone matrix, resulting in aberrant vascularisation and osteogenesis.^10^ Hypertrophic chondrocytes also participate in this process as the major source of VEGF, coupled vessel invasion, cartilage remodelling, and ossification.^11^ Interestingly, ECs recruited by multiple biological agents are the major driving forces of cartilage matrix degradation, bone elongation, and remodelling.^12^ Moreover, OB-derived VEGF participates in the delicate bone-vessel crosstalk,^13^ however, the specific communication mode inside the OA subchondral bone remains unknown. OBs have multiple functions, including angiogenesis promotion, matrix manufacturing and mineralisation.^14^ Simultaneously, the participation of various immune cells leads to aggravated inflammation and subchondral bone disorders.^15^ This indicates the importance of subchondral bone cells in OA progression, and the multifaceted nature of ECs and OBs suggests that they are comprised of diverse subpopulations. Nevertheless, considering the high phenotypic heterogeneity and limited understanding of biomarkers, the isolation and definition of EC and OB subtypes in human subchondral bone remains unclear.

To reveal the cellular interactions involved in the OA subchondral environment, single-cell RNA sequencing (scRNA-seq) was performed on tibial subchondral bone samples from patients undergoing total knee arthroplasty. Here, the scRNA-seq technique was utilised to map a general census of subchondral bone cells from both normal and OA sites, determine the genetic characteristics of these cell subgroups, and further analyse their potential differentiation relationships to characterise specific cell types. Finally, we investigated the cell-cell interaction network between EC and OB subpopulations. These results expand our understanding of the heterogeneity between patients and provide a theoretical basis for personalised OA therapies.

## RESULTS

### Single-cell profiling of human OA subchondral bone cells

To identify the cellular constitution of subchondral bone cells in human OA, we isolated human OA subchondral bone cells obtained from both lateral and medial tibial plateaus of two patients undergoing knee arthroplasty and profiled subchondral bone cells from different locations and patients (n=4) using scRNA-seq (Figure1A and supplementary Table S1). Through unbiased clustering of human subchondral bone cells, we found 10 clusters from OA patients were identified, including T (11151), B (990), NK (5155), NKT (7538), and dendritic cells (DCs; 124), monocytes and macrophages (557), bone-related cells (864): ECs (246), mesenchymal stem cells (MSCs; 104), and OBs (360). According to animal experiments and clinical experience, OA involvement is more common and occurs earlier in the medial side of the tibial plateau. In this study, we selected patients with severe medial destruction and an almost healthy lateral plateau. These two patients were fully informed of their condition and chose total knee arthroplasty. Subchondral bone cells were divided into the control group (Ctrl; 13024) from the lateral tibial plateau and the OA group (13355) from the medial side (Figure1B). In total, 26379 cells were retained for subsequent analysis after rigorous filtration (Figure1C, D; Supplementary FigureS1A, S2A and supplementary Table S2).

**Figure1.**
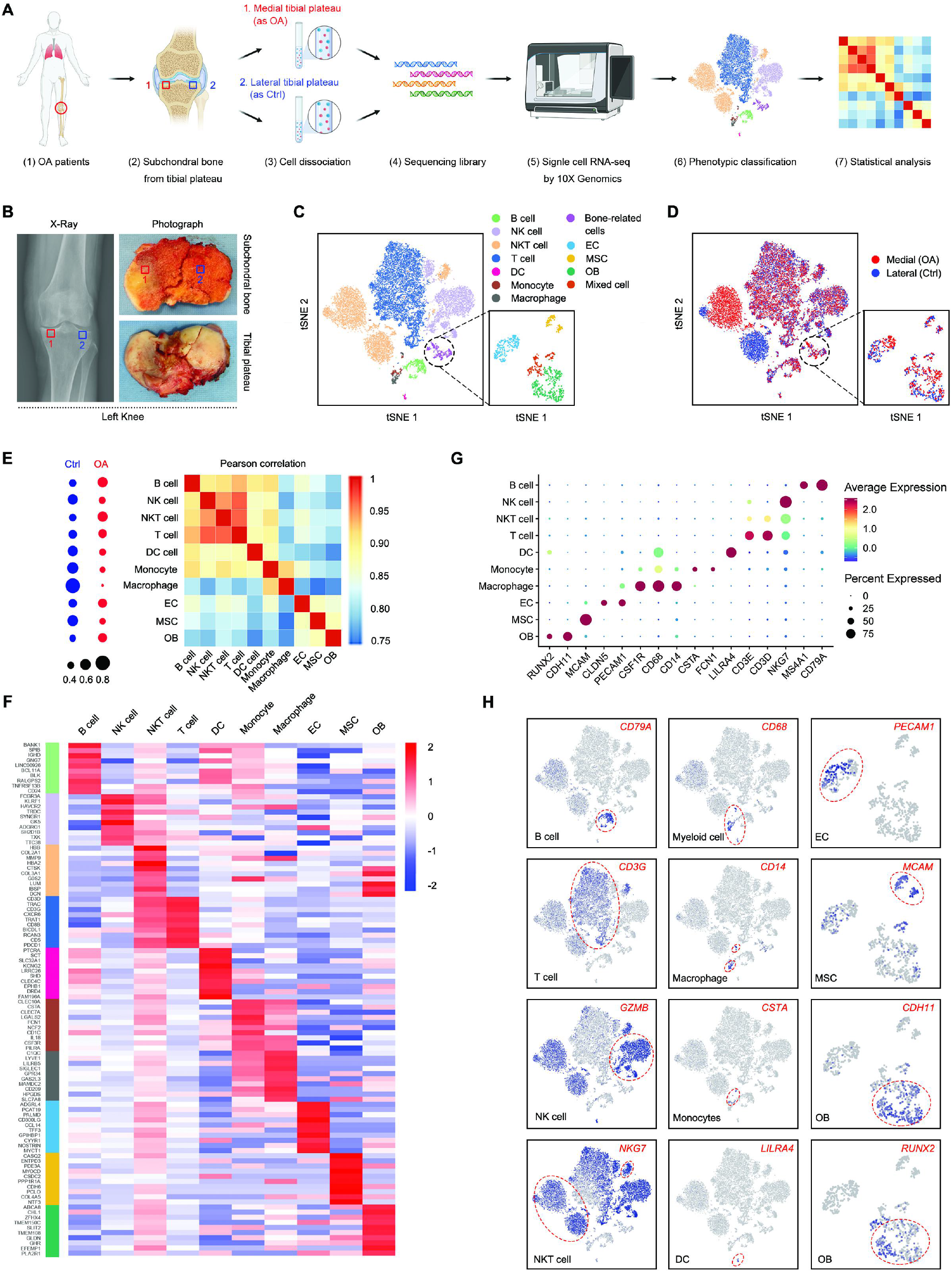
Single-cell profiling of human OA subchondral bone cells. (A) Schematic workflow of the experimental strategy. (B) X-ray photograph of a patient with knee osteoarthritis (OA; left) and corresponding cross-sectional anatomy of the subchondral bone and tibial plateau (right). (C,D)The tSNE plots (left panel) and the sample origin (right panel) of 26,379 subchondral bone cells and 864 bone-associated cells. (E) Dot plots showing the distribution of each cell type in the control (Ctrl) and OA groups. Heatmap showing the pairwise correlations. (F) Cluster averaged log-normalised expression of the top 10 marker genes between the 10 cell types with stromal-related genes of interest annotated. Expression values are scaled per cluster. (G) Dot plot showing the expression of specific signatures in identified cell types in (F). The dot colour and size represent the mean expression and proportion of each cell population expressing genes, respectively. (H) Feature plots showing the expression of indicated markers for each cell type on the t-SNE map.

Next, the cell type distribution in the Ctrl and OA groups was analysed. In addition to the relationship between immune and myeloid cells, paired correlation analysis showed tight connections between ECs and MSCs, ECs and OBs, and MSCs and OBs (Figure1E). The first ten upregulated genes from these ten clusters were used to create a heatmap (Figure1F). Representative markers for T, B, NK, and NKT cells, DCs, monocytes, macrophages, ECs, MSCs, and OBs were revealed (Figure1G). Specifically, the following clusters were identified: (1) B cells (expressing CD79A, BANK1, and MS4A1), (2) NK cells (expressing GZMB and NKG7), (3) NKT cells (expressing NKG7 and CD3D), (4) T cells (expressing CD3D and CD3G), (5) DCs (expressing LILRA4 and PTCRA), (6) monocytes (expressing CSTA and FCN1), (7) macrophages (expressing CD14, CD68, CSF1R, C1QC, and F13A1), (8) ECs (expressing PECAM1 and CLDN5), (9) MSCs (expressing MCAM), and (10) OBs (expressing RUNX2 and cadherin 11 [CDH11]; Figure1H and Supplementary FigureS1B, S2B–D).

### Identification of bone-related cell populations in human OA subchondral bone

Abnormal angiogenesis, subchondral bone remodeling and sensory innervation are well recognized during early stage of OA, and might cause cartilage destruction and pain directly or indirectly.^13^ Therefore, OBs, osteoclasts, ECs, and neuronal cells were the focus of our research, rather than immune cells. To define the cell subpopulation and identify genome-wide gene expression patterns, bone-related cells were clustered to produce eight clusters (Figure2A). Next, to explore the potential transformation between different cell types and visually depict the differentiation paths, the Monocle method was used to determine the pseudotemporal order between cell types (Figure2B, C). Our analysis clearly identified and verified three major groups of differentiated cell types: ECs (PECAM1+), MSCs (MCAM+), and OBs (RUNX2+/CDH11+; Figure2D).

**Figure2.**
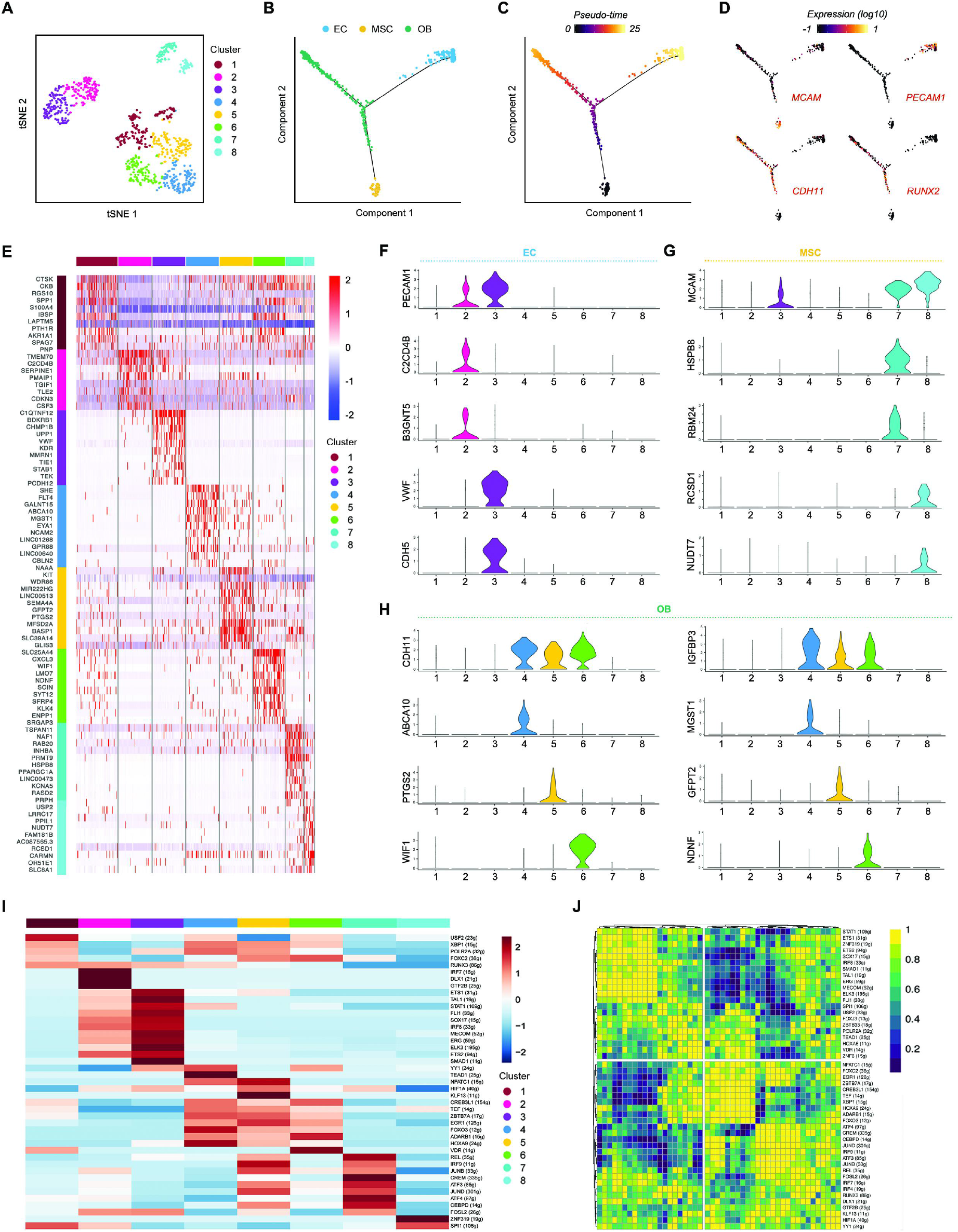
Identification of bone related cell populations in human OA subchondral bone. (A) t-SNE plots of bone associated cells coloured by cluster. (B–D) Pseudotime trajectory plot showing differentiated cell types (endothelial cells [ECs], mesenchymal stem cells, and osteoblasts [OBs]) at the end of the branches. Dots along the trajectory lines represent the status of the cells transitioning toward differentiated cell types. (E) Heatmap revealing the scaled expression of differentially expressed genes (DEGs) for each cluster defined in (A). (F–H) Violin plots showing expression levels of indicated markers for eight clusters. (I–J) Single-cell regulatory network inference and clustering analysis showing distinct regulons in eight clusters. The heatmap shows only the regulons with significant differences.

Among the eight clusters, cluster1 expressed markers of multiple cell types, such as CTSK, RGS10, and SPP1, suggesting that cluster1 contains osteoclasts, nerve cells, and OBs (Supplementary Figure3). Cluster1 is a heterogeneous cell cluster; therefore, we only analysed OBs, ECs, and MSCs. To determine the characteristics of each cell cluster by analysing differential gene transcript expression patterns, a differentially expressed feature analysis was performed using the scRNA-seq dataset and all cell clusters were compared with one another. We discovered 78 differentially expressed genes (DEGs) that best divided bone-related cells in subchondral bone into eight subclusters (Figure2E). Next, the original sample information and expression levels of the indicated markers were combined to determine the cell identity of each cluster, and their biological functions were analysed via regulons CSI correlation heatmap of the co-expression between transcription factors (TFs) and potential target genes. Seven major cell clusters were identified: precursor ECs, pre-ECs (C2CD4B+/B3GNT5+); ECs (VWF+/KDR+); endothelial OBs, EnOBs (ABCA10+/microsomal glutathione S-transferase 1 [MGST1]+); stromal OBs, StOBs (PTGS2+/ glutamine-fructose-6-phosphate transaminase 2 [GFPT2]+); mineralised OBs, MinOBs (WNT inhibitory factor 1 [WIF1]+/NDNF+), and two MSC subpopulations (Figure2F–H). Through SCENIC analysis, we discovered that pre-ECs and ECs exhibit activated pro-angiogenesis regulons, such as SMAD1, ERG, and ETS1, and that the regulators have higher activity in ECs than in pre-ECs. Furthermore, OB subpopulations exhibited similar activated TFs, however, the regulatory activities of TFs are different (Figure2I, J).

### Identification of pre-ECs and ECs

In addition to the well-known differences between arteries, capillaries, and veins, ECs are highly heterogeneous and acquire specialised functional properties in the local microenvironment. The articular cartilage is constantly maintained in a low-oxygen environment. However, due to the high metabolic requirements of OBs, blood vessels are required to provide sufficient oxygen. The cells are relatively hypoxic during osteogenesis. Osteogenesis and nearby ECs may increase HIF-1α expression. Upregulation of HIF-1α activity in hypoxic tissues leads to increased VEGF expression and promotes angiogenesis. Using immunofluorescence and flow cytometry to identify the EC subpopulation in bone, Kusumbe et al. proposed the following terminology for bone microvessels: H-type for the small PECAM1^hi^/Emcn^hi^ subset and L-type for the PECAM1^lo^/Emcn^lo^ sinusoidal vessels.^16^ As previously mentioned, EC subpopulations were divided into pre-ECs and ECs, and we discovered certain differences between them (Figure2E, F). To investigate the distinct features of pre-ECs and ECs, we identified DEGs between them (Figure3A). The Gene Ontology (GO) and Kyoto Encyclopaedia of Genes and Genomes (KEGG) were analysed with these DEGs to show pre-EC and EC characteristics. Notably, pre-ECs were enriched for extracellular exosomes, interleukin-mediated signalling pathways, and ribosomes, whereas ECs were enriched for vasculogenesis, angiogenesis, EC migration, and signalling pathway regulation, such as VEGF, Rap1, PI3K/Akt, Ras, and MAPK signalling pathways (Supplementary FigureS4A–D). Then we compared characteristic genes of these two clusters Pre-EC identification markers have diverse functions, including genes related to ribosome synthesis, exosome synthesis, and inflammation, including RPL17, HNRNPF, RABA5, and C2CD4B, whereas ECs are primarily enriched in genes related to angiogenesis, such as VWF, KDR, TIE1, and CDH5 (Figure3B). Moreover, angiogenesis-related EMCN, PECAM1, EGFL7, ENG, and KDR were all upregulated during the differentiation of pre-ECs to ECs, while exocrine-related DDIT3 and RAB5A73 and inflammation-related CCL2 and C2CD4B74 were downregulated (Figure3C).

**Figure3.**
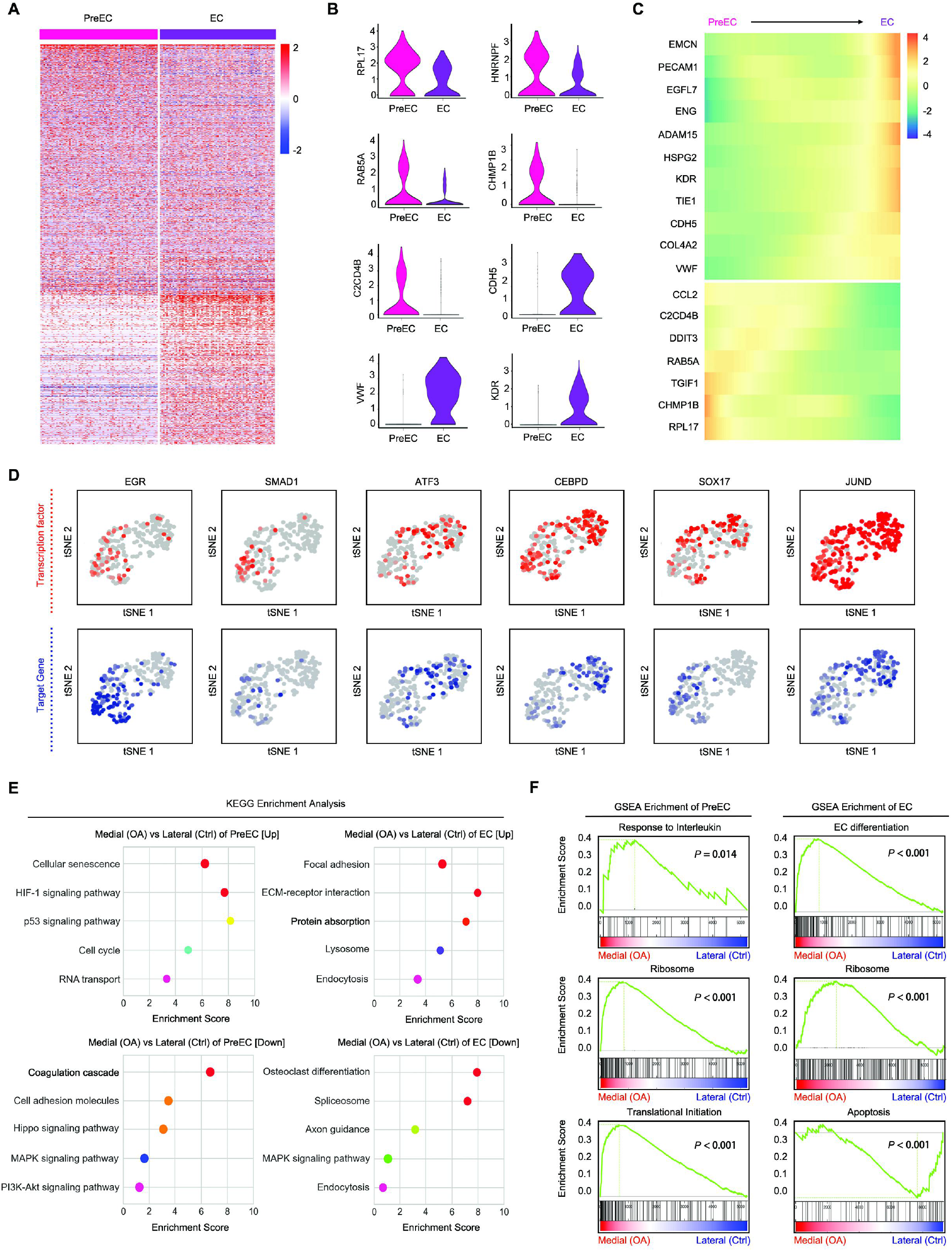
Identification of precursor ECs (pre-ECs) and ECs. (A) Heatmap of DEGs between different ECs. (B) Violin plots showing the expression levels of the specific representative genes marking pre-ECs and ECs. (C) Heatmap showing upregulation or downregulation of vascular markers, exosomes, and ribosomal markers in the differentiation process. (D) tSNE plots of the expression levels of transcription factors (TFs; up) and area under the curve scores (down). (E) Kyoto Encyclopaedia of Genes and Genomes pathway enrichment between the OA and Ctrl groups in pre-ECs and ECs. (F) Gene Set Enrichment Analysis (GSEA) pathway enrichment between the OA and Ctrl groups in pre-ECs and ECs.

Next, the essential motifs of the two EC subpopulations were identified using SCENIC analysis. As the specific motifs of pre-ECs, ATF3 and CEBPD are essential in the transcriptional regulation of inflammation, whereas ERG and SMAD1 motifs, which are closely related to angiogenesis, are highly activated in ECs. SOX17 and JunD are TFs that are widely expressed in pre-ECs and ECs (Figure3D and Supplementary FigureS5A,B). In a previous study, SMAD1 sprouted angiogenesis in human embryonic stem cell-derived ECs.^17^ ERG is an essential regulator of angiogenesis and vascular stability through Wnt signalling.^18^ These results may help us to identify pre-ECs and ECs and expand our understanding of the novel function of subchondral EC subsets in OA.

During OA progression, the EC cluster was a substantially increased cell population (Figure1E). To investigate the distinct features of the OA and Ctrl groups, we identified DEGs between these two groups in pre-ECs and ECs (Supplementary FigureS6A,B). To analyse the identified features of pre-EC and EC clusters from the OA group, we analysed the differences using GO, KEGG, and Gene Set Enrichment analyses (GSEA). Notably, compared with that of the Ctrl group, pre-ECs of the OA group showed stronger protein synthesis, inflammation-related pathways, and responses, whereas ECs were enriched for angiogenesis-promoting functions, such as blood vessel development, EC differentiation, and platelet-derived growth factor binding (Figure3E,F and Supplementary FigureS6C,D).

### Determining the relationships among EnOBs, StOBs, and MinOBs

According to Rutkovskiy et al., OBs undergo a 3-stage differentiation: Stage 1, the cells continue to proliferate; Stage 2, they start differentiating, while maturing the extracellular matrix (ECM) with alkaline phosphatase (ALP) and collagen; and Stage 3, the matrix mineralisation and mineral deposits increase.^19^ According to the SCENIC analysis (Figure2I), we found that the TF types and transcriptional activities were different among EnOBs, StOBs, and MinOBs, which means that these clusters may have different cellular functions. By identifying DEGs between EnOBs, StOBs, and MinOBs, the differences between OB subpopulations were further verified (Figure4A). To explain the specific characteristics of these three OB populations, we analysed the differences through GO and KEGG analyses. EnOBs are related to EC migration, VEGF binding, and the PDGFR-β signalling pathway, and express NRP1, PDGFRB, and VCAM, suggesting that this cluster may have potentially affect angiogenesis. StOBs were enriched for collagen and fiber-related biological processes, such as collagen fibril organisation, fibronectin binding, and ECM binding. MinOBs distinctively expressed an ossification and bone mineralisation biological process gene signature (Figure4B and Supplementary FigureS7A–C). Next, representative candidate markers among EnOBs, StOBs, and MinOBs (Figure4C) were identified, including TFs (Figure4F).

**Figure4.**
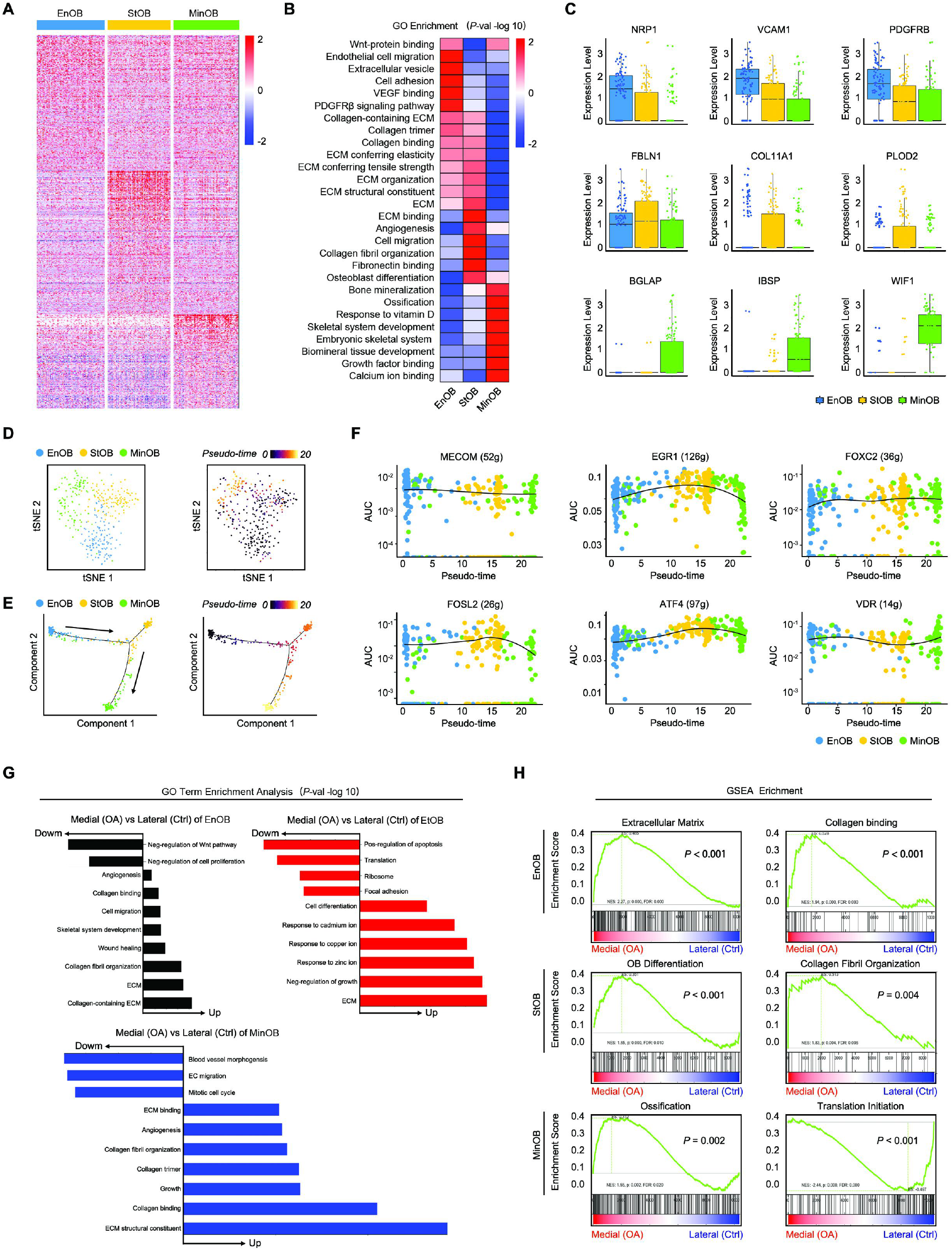
Determining the relationships among endothelial OBs (EnOBs), stromal OBs (StOBs), and mineralised OBs (MinOBs) (A) Heatmap showing Z score scaled expression levels of DEGs for EnOB, StOB, and MinOB populations. (B) Heatmap showing the differences in enriched Gene Ontology (GO) functions of upregulated genes in different OB subsets. (C) Boxplots showing the expression levels of representative candidate marker genes specifically expressed in different subsets. (D,E) Monocle pseudospace trajectory revealing the progression of OB lineage in subchondral bone coloured according to cluster. Monocle pseudotime trajectory revealing the progression of EnOBs, StOBs, and MinOBs. (F) Pseudotemporal expression dynamics of TFs in EnOBs, StOBs, and MinOBs. All single cells in the EnOB, StOB, and MinOB cell lineage are ordered based on pseudotime. (G) GO functions enrichment analysis of OA vs. Ctrl upregulated genes in EnOBs, StOBs, and MinOBs. (H) GSEA showing enrichment of pathways between the OA and Ctrl groups in EnOBs, StOBs, and MinOBs.

Next, the Monocle method was applied to depict the pseudotemporal sequence of potential differentiation pathways among cell types. According to the pseudotime trajectory axis, we suggest that StOB is an intermediate state representing the state between EnOBs and MinOBs (Figure4D,E). The pseudotemporal expression dynamics of representative candidate markers and TFs also marked the progression from EnOBs to StOBs to MinOBs. Through SCENIC analysis, we found that TFs upregulated by EnOBs were related to angiogenesis, such as MECOM, ERG, and XBP1, and TFs related to collagen and fibre in StOBs, including FOSL2, ERG1, and ATF4, and VDR and FOXC2 in MinOBs are related to mineralisation (Figure4F and Supplementary FigureS8A–C). Taken together, these data reveal the relationships and potential functions of EnOBs, StOBs, and MinOBs.

The number of OBs increased in abundance during OA progression, more so than in ECs (Figure1E). To investigate the distinct features of normal and diseased cells inside the subchondral zone, we identified DEGs between the OA and Ctrl groups of these three OB clusters, respectively (Supplementary FigureS9A–C). We then analysed the differences by GSEA and GO analysis. Notably, compared with that of the Ctrl group, the OA group of EnOBs showed stronger angiogenesis and wound healing biological processes; the OA group of StOBs was enriched for ECM binding and collagen fibril organisation, and the OA group of MinOBs was enriched for response of metal ions such as cadmium, copper, and zinc (Figure4G,H). Regarding these ionic reactions, copper ions in biological materials promote bone formation,^20^ zinc increases OB activity and collagen synthesis,^21^ and cadmium promotes OB differentiation.^22^ These results indicate that OBs in the OA group have stronger osteogenic effects and we will further verify the functional heterogeneity of the three OB subpopulations.

### Vascular EC and OB subpopulation interaction

Through the interaction of H-type blood vessels and various cytokines in bone metabolism, angiogenesis and bone formation are precisely coupled.^16^ MSCs are chemically recruited by ECs to promote osteogenesis.^23^ Furthermore, the crucial signalling pathways in MSCs coupled with ECs include the TGF-β, PDGF-PDGFR, angiopoietin, Notch, and FAK signalling pathways. To further explore the key signalling pathway that couples OBs and ECs, we studied the cell-cell interaction network between the identified clusters. Considering the outcomes of GSEA and GO enrichment analyses and the characteristics of the genes and TFs, we chose to analyse the interaction pairs, including chemokines, ephrin receptor family, NOTCH family, cytokines, and integrins. We found that ECs were the predominant cell population interacting with the OB subpopulation, and pairwise correlation analysis revealed that ECs are more closely related than pre-ECs to OB subpopulations (Figure5A, B).

**Figure 5.**
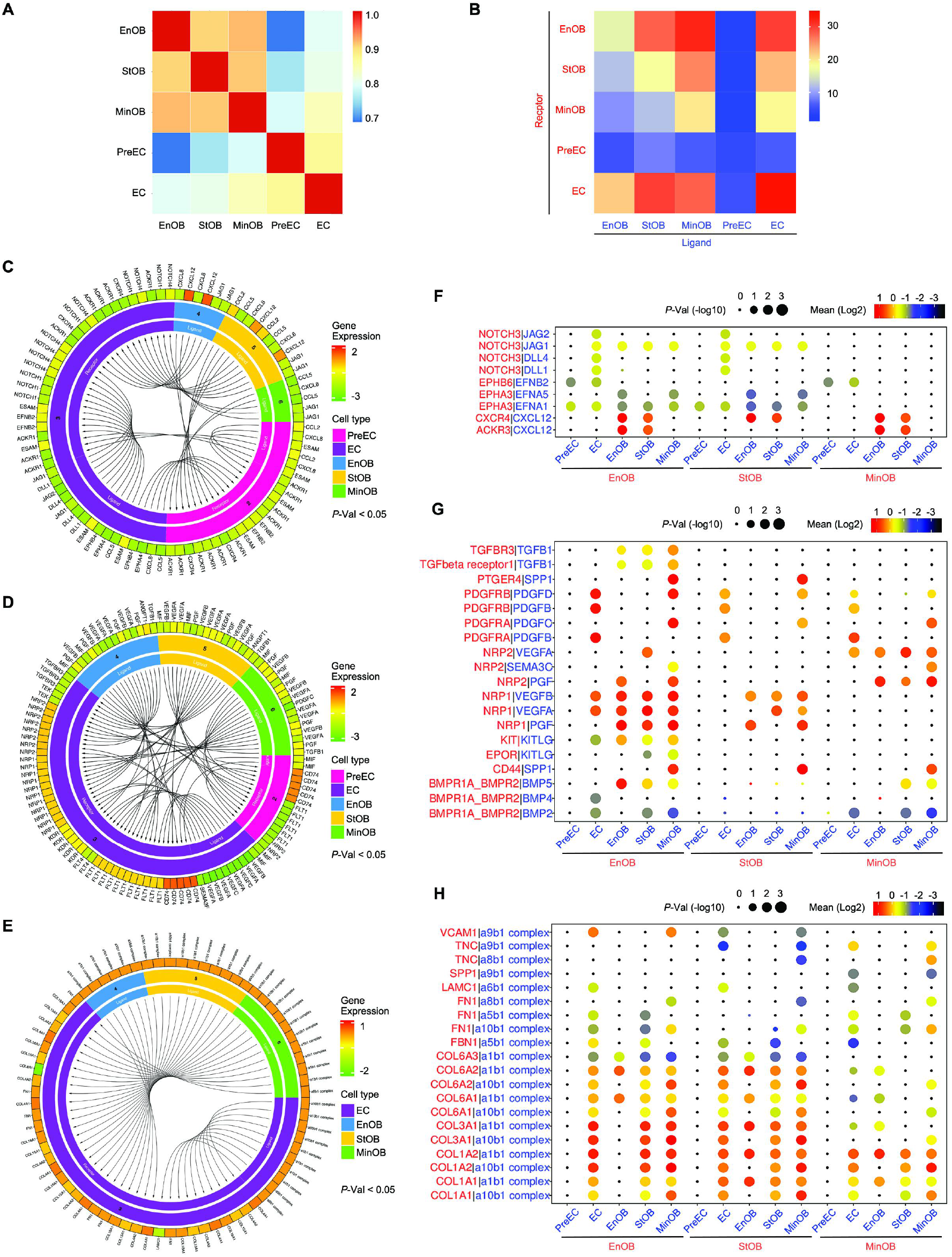
Vascular EC and OB subtype interaction. (A) Pearson correlation analysis of two clusters of ECs subsets and three clusters of OBs. (B) CellPhoneDB analysis showing the number of ligand-receptor interactions between EC and OB subpopulations. Circos plots showing ligand-receptor pairs of cytokines (C), growth factors (D), and integrin (E) between EC subpopulations. Bubble plots showing ligand-receptor pairs of cytokines (F), growth factors (G), and integrin (H) between OB subpopulations.

Compared with pre-ECs, ECs express a larger number of membrane receptors, fibronectin and collagen, and secrete a larger number of angiocrine factors. EC analysis revealed that ECs exhibited abundant expression of multiple membrane receptors for ligands important for vascular development, including NOTCH1, NOTCH4, VEGF receptors (KDR, FLT1, FLT4, NRP1, NRP2), TGFβ receptors (TGFBR3), ephrin B receptor (EPHB4), and tyrosine kinase receptor (TEK), which bind to JAG1, the VEGF family, PGF, ANGPT1, and TGFβ 1 ligand secreted by OB to promote angiogenesis (Figure5C–E and Supplementary FigureS10A–C). These results indicate that ECs are a mature EC subgroup with angiogenic function at the transcriptional level, and are mainly coupled with OBs. Notably, pre-ECs hardly secrete any ligands, as the results showed, however, considering the enrichment analysis data of pre-ECs (Supplementary FigureS4A), we hypothesise that the function of pre-ECs is achieved by secreting exosomes, which requires further investigation.

There is little difference between chemokine interaction pairs in OBs, and they all express CXCR4 to promote OB proliferation and differentiation, however, only MinOBs do not secrete CXCL12.^24^ At the end of osteogenic differentiation, CXCL12 is downregulated.^25^ We found that the overall expression of the ephrin receptor and NOTCH family members decreased gradually from EnOBs to StOBs to MinOBs. The ephrin receptor^26^ and NOTCH families^27^ promote OB proliferation, and the lack of the bone system JAG1 leads to mature OB proliferation, which is manifested as an increase in the rate of mineral deposition (Figure5F). Additionally, we found that these three OB subgroups are affected by bone morphogenetic proteins, which have a strong positive effect on bone formation.^28^ PGF, PDGF, and VEGF families secreted by ECs and OBs could interact with the receptors highly expressed on the surface of EnOBs, including NRP1, PDGFRA, and PDGFRB. These cytokines are related to angiogenesis, and PDGF induces OB proliferation via the ERK signalling pathway.^29^ We also found that TGFβ1 was produced by all three types of OBs, but only bound with TGFβ receptors on EnOBs to play a role in promoting OB proliferation,^30^ inducing VEGF secretion^31^ and inhibiting mineralisation function.^32^ As a non-collagenous protein in the bone ECM that is recognised to regulate bone formation and mineralisation, osteopontin (OPN/SPP1) is expressed and released in the integrin interaction pair by MinOBs only (Figure5G, H). Compared with that of EnOBs and MinOBs, StOBs expressed the highest fibrin and collagen levels, while MinOBs expressed the lowest (Figure5H). In summary, the above data further validated our understanding of the role of these five major cell clusters in bone-associated cells and the interaction between ECs and OBs.

### Pathological identification of subpopulations

In order to further verify our sequence results, we conducted destabilisation of the medial meniscus (DMM) surgery on 6-week-old C57B6J mice, simulating patient conditions before arthroplasty surgery (Figure6A). The tibial subchondral bone volume in OA mice showed significant changes after surgery, as shown by microCT, safranin O, fast green staining, and H&E staining (Figure6B). The total tissue volume of subchondral bone decreased at 2 weeks and increased at 4 weeks post-surgery, and the subchondral bone structure densified at 8 weeks (Figure6C). Microstructure disruption was indicated by aberrant subchondral bone plate thickness and trabecular pattern factor, according to the microCT calculation (Figure6C). Osteoarthritis Research Society International scores indicated that cartilage degeneration was significant at 4 weeks and deteriorated at 8 weeks post-surgery (Figure6D).

**Figure 6.**
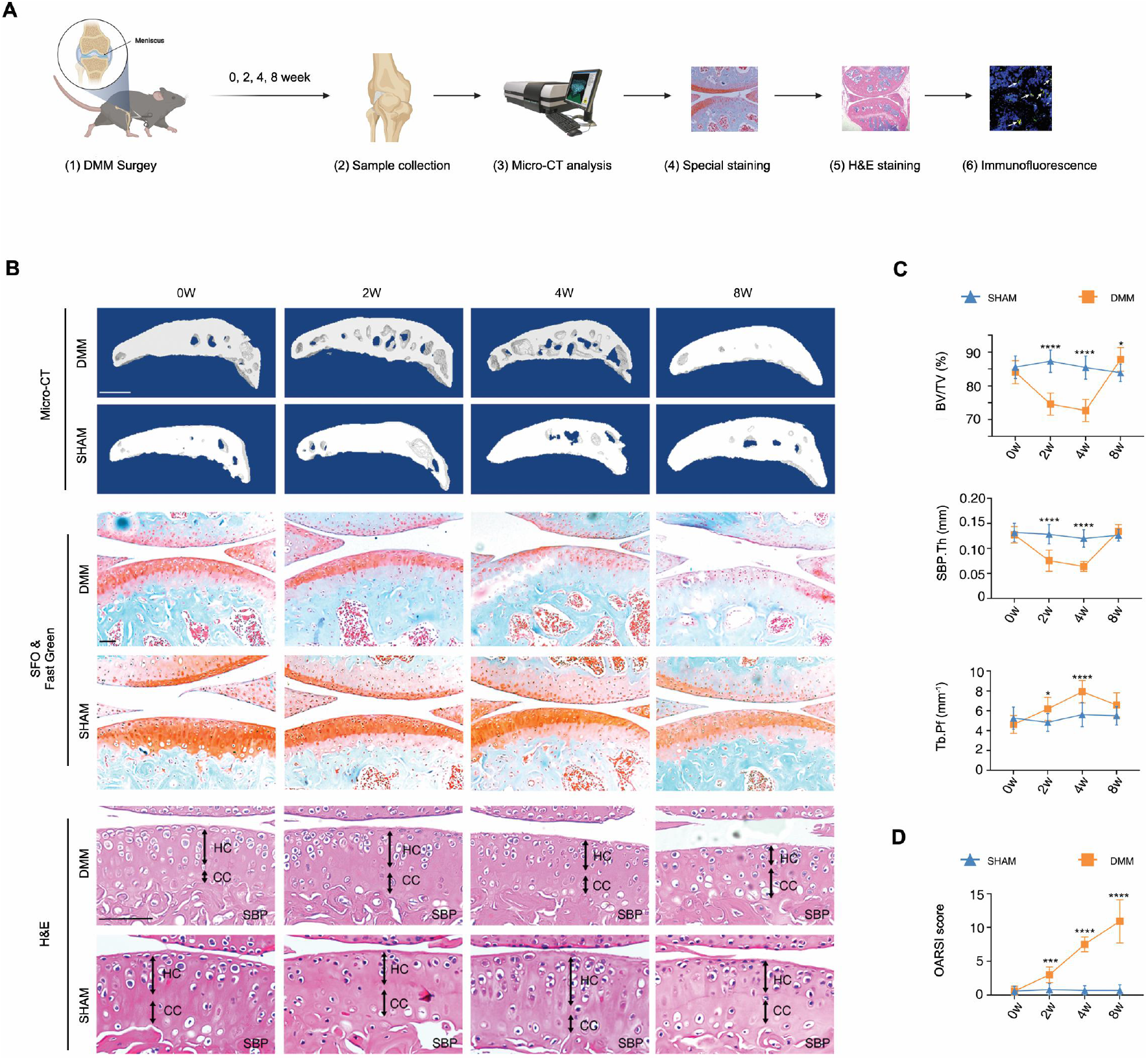
Pathological identification of DMM mice model. (A) Schematic illustration of the experimental process. (B) Top row, representative 3-dimension image of subchondral bone from tibial medial plateau at 0, 2, 4, and 8 weeks after sham or DMM surgery. Middle rows, safranin O and fast green stain of tibial medial plateau at the same checkpoint. Bottom rows, H&E staining of knee joint, cartilage and subchondral bone. HC: hyaline cartilage; CC: calcified cartilage; SBP: subchondral bone plate. (C) Quantitative analysis of bone volume tissue/total tissue volume (BV/TV), subchondral bone plate thickness (SBP. Th) and trabecular bone pattern factor (Tb.Pf) in medial plateau calculated from micro-CT results. (D) Osteoarthritis Research Society International scores after DMM operation are shown bottom right. n=10 in each group. Scale bar, 100 μm. \**p*<0.05, \*\**p*<0.01, \*\*\**p*<0.001, \*\*\*\**p*<0.0001. Statistical significance was shown by Two-way ANOVA. Data was presented as the mean±SD.

An immunofluorescence test was performed to characterise cell development and trajectory during OA pathology. The ratio of ECs among the total PECAM1-positive cells increased continuously after surgery (Figure7A). Specifically, PECAM1-positive ECs were rather rare and were near the trabecular before intervention, indicating a relatively low angiogenesis rate. An important proportion of PECAM1-positive cells were KDR-negative, or pre-ECs, in the sham group (66.4±9.44%) and 0w samples in the DMM group (60.69±8.77%). They are characterised by genes coded for ribosome synthesis, extracellular vesicles synthesis, and inflammation, indicating hypermetabolism and pro-inflammatory status inside the subchondral bone. ECs grew more after 4 weeks and the ratio of KDR-positive, or ECs with relatively high angiogenesis trends, increased significantly and reached 82.97±8.01%. With severe subchondral sclerosis progression, the majority (88.90±4.62%) of ECs became pre-ECs, and the total number of pre-ECs and ECs decreased dramatically in the limited space (Figure7B).

**Figure 7.**
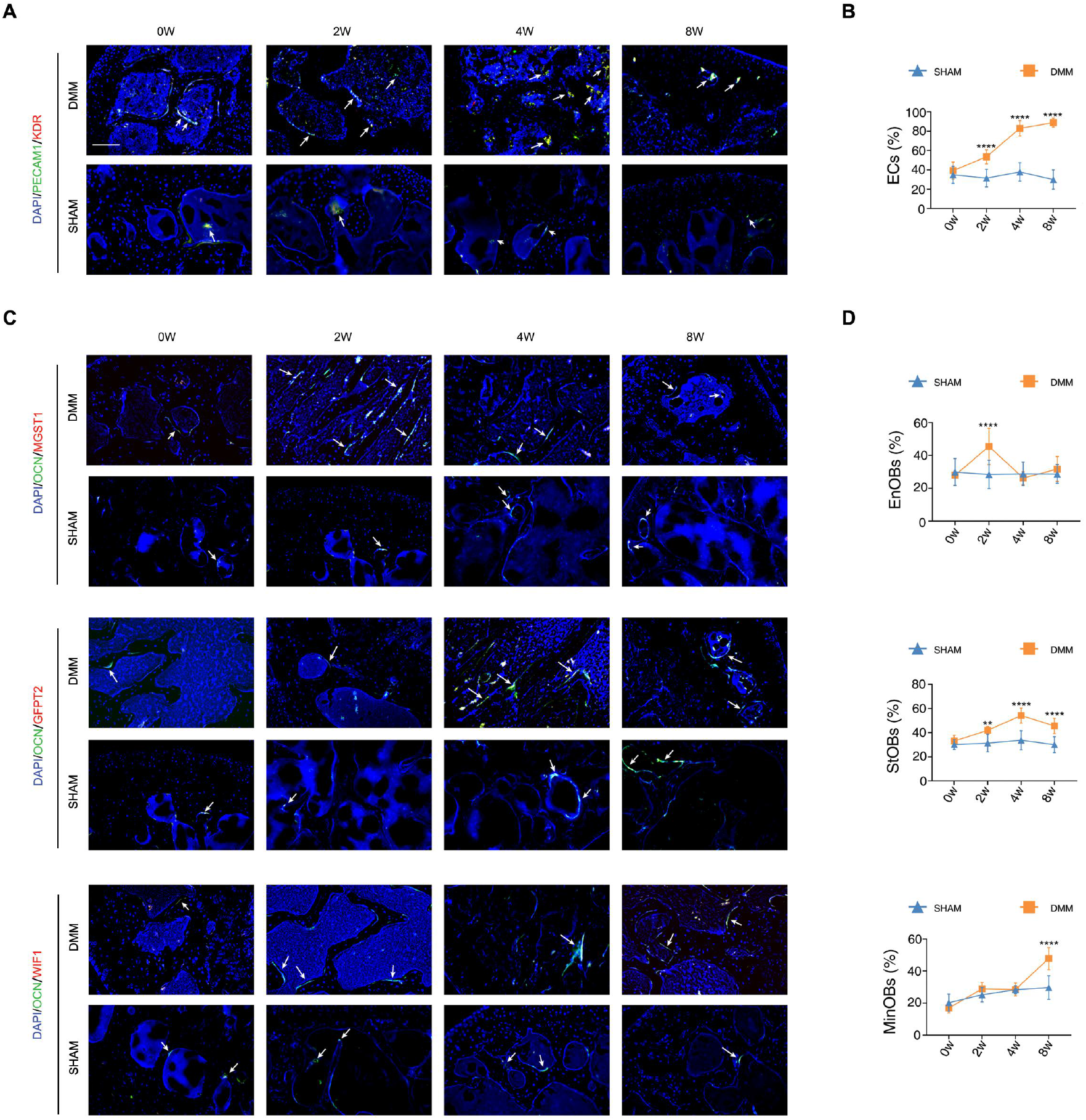
Pathological identification of EC and OB subpopulations. (A) Immunofluorescence staining of PECAM1, KDR, white arrow shows co-positive area of corresponding markers. (B) Percentage of PECAM1+KDR+ ECs in total PECAM1+ cells. (C) Immunofluorescence staining of OCN, MGST1, GFPT2 and WIF1, white arrow shows co-positive area of corresponding markers. (D) Percentage of MGST1+EnOBs, GFPT2+ StOBs and WIF1+ MinOBs in OCN+ cells, n=9 in each group. Scale bar, 100 μm. \**p*<0.05, \*\**p*<0.01, \*\*\**p*<0.001, \*\*\*\**p*<0.0001. Statistical significance was shown by Two-way ANOVA. Data was 652 presented as the mean±SD.

Next, we analysed the OB subpopulation marked by osteocalcin (OCN), MGST1, GFPT2, and WIF1. Consistent with previous results, the total number of OCN-positive cells increased during the first 4 weeks and decreased at 8 weeks post-surgery (Figure7C). Analogously, EnOBs characterised by angiogenesis-related genes increased dramatically 2 weeks post-DMM surgery and decreased at 8 weeks (Figure7D, top). StOBs capable of ECM binding and collagen fibril organisation increased continuously during the first 4 weeks post-trauma and decreased at 8 weeks (Figure7D, middle), probably because of the calcification requirement. MinOBs, which were closely related to metal ion and biomineralisation, increased dramatically at 8 weeks post-surgery and accounted for approximately 48% of the total OCN-positive cells (Figure7D, bottom). Taken together, these data indicate chronological changes in OB subgroups and mapping the over-time changing bio-function of OBs in subchondral bone during post-traumatic OA.

## Discussion

OA is one of the most common degenerative diseases that cause disability in older adults. An epidemiological study by Tang et al. showed that 8.1% of the adult population had clinically significant OA of the knee or hip.^33^ Moreover, OA consumes a substantial amount of healthcare resources, primarily owing to the joint replacement surgery costs for advanced OA.^33^ Increasing economic pressure, an aging society, and the obesity epidemic emphasise the need for new strategies for the diagnosis and intervention at early-stage OA.^34–36^ Increasing evidence suggests that the appearance of subchondral bone lesions occurs earlier than cartilage degeneration, and the pathological alterations of subchondral bone play an important role in OA development. During the initial phase of OA, the bone turnover rate beneath the articular cartilage was upregulated, and vascular invasion took place bottom-up from the subchondral, the tidemark, and ultimately into the cartilage. Since the exact role of subchondral bone during OA initiation and progression remains unclear, and the specific cell markers are lacking, it is urgent to unveil the internal state of subchondral bone cells in OA pathogenesis. Here, we used comprehensive gene expression profiling to reveal the cell types that make up the subchondral bone microenvironment at single-cell resolution, and also novel cell markers and characteristics to verify each hypothetical subchondral bone cell cluster.

We identified 10 different cell types in the human OA subchondral bone microenvironment. Notably, there were more OBs and ECs in the OA group than in the Ctrl group. This finding validates the characteristics of increased bone formation and angiogenesis in OA subchondral bone.^37^ In addition to the empirically inferred subchondral bone cell types, we identified new subtypes of bone-associated cells and new markers of bone-associated cell populations based on scRNA-seq analysis. Based on the expression of TFs and markers of the new subtypes, we suggest that EC subtypes and OB subtypes perform different biological functions.

In the process of OA cartilage erosion, compared with the top-down vessel invasion originating from synovial tissue or synovium, bottom-up vascularization from subchondral bone plays a larger role.^38^ In the process of bone elongation, ECs constantly erode the cartilage matrix, thus creating space for osteogenesis. Kusumbe et al. identified two types of special blood vessel subtypes based on the expression strength of PECAM1 and EMCN, namely H-type (PECAM1^hi^EMCN^hi^) and L-type blood vessels (PECAM1^lo^EMCN^lo^).^16^ In the present study, we also identified two EC phenotypes: pre-ECs and ECs. We found that PECAM1 and EMCN expression in ECs was upregulated, suggesting that it may have functions similar to those of H-type blood vessels. By comparing gene expression, TF activity, and enrichment analysis, we suggest that the “ECs” in this research are a type of endothelial cells that promote angiogenesis, and can also be coupled with OBs. In this study, we identified a new subset named pre-ECs, characterised by interleukin-mediated inflammation pathways and exosomes. A key element leading to the advancement of OA is the production of high inflammatory cytokine levels, and the pro-inflammatory cytokine interleukin 1β (IL-1β) is expressed in large quantities during OA. Elevated IL-1β levels are associated with tissue damage and reflect the severity of inflammation. During subchondral bone reconstruction, IL-1β may also play a role in promoting cartilage calcification (ossification) and cartilage degeneration.^39^ Yang et al. showed that exosomes derived from vascular ECs promoted the progression of OA by promoting chondrocyte apoptosis.^40^ Therefore, we speculate that pre-ECs in OA may allow for bone formation through exosomes digesting cartilage and promote subchondral bone remodelling through IL-mediated inflammation. The characterisation of these two novel subsets improves our understanding of the characteristics and functions of ECs in the OA subchondral bone.

Subchondral bone sclerosis is characterised by an increase in bone volume due to an enhanced bone turnover rate.^37^ In the OA subchondral bone, we found three OB phenotypes: EnOBs, StOBs, and MinOBs. We believe that EnOBs are enriched in angiogenesis-related pathways, StOBs are characterised by collagen and fibrosis, and MinOBs specifically express mineralization-related markers. Here, we found that the three OB phenotypes have a differentiation path, from EnOBs to StOBs to MinOBs. The first stage of OB differentiation is characterised by cell proliferation. EnOBs are tightly correlated with vascularisation, including EC migration, VEGF binding, and platelet-derived growth factor receptor β (PDGFRβ) signalling pathways. Angiogenesis directly promotes bone formation, and CDH11 cooperates with PDGFRβ to promote cell proliferation. In the next stage, OBs begin to differentiate and express collagen and alkaline phosphatase ALP. StOBs are OBs characterised by OB differentiation and collagen fibril organisation, and express high levels of fibulin-1 (FBLN1), collagen type XI alpha 1 (COL11A1), and PLOD2. FBLN1 is an important ECM protein that stabilises collagen and other ECM proteins. COL11A1, as one of the three alpha chains of type XI collagen, is critical for collagen fiber assembly and bone development. OA-related fibrosis is associated with elevated PLOD2 expression.^41^ In the last stage, OBs express markers of more bone sialoprotein, OPN/SPP1, and osteocalcin (BGLAP), thereby inducing matrix mineralisation. In this study, we found that MinOBs specifically express the above-mentioned mature OB phenotypic markers, representing the biological processes of ossification, bone mineralisation, and biomineral tissue development. Additionally, MinOBs express the Wnt antagonist WIF1, which regulates the Wnt/β-catenin signalling pathway to reduce cell proliferation and promote mineralisation.^42^

The coupling of osteogenesis and angiogenesis causes increased bone mineral density and significant microstructural changes in the subchondral bone. Through the analysis of cell-to-cell communication between ECs and OBs, we found that ECs with angiogenic function and upregulated PECAM1 and EMCN expression are coupled with OBs through NOTCH, VEGF, and TGFβ receptors, EPHB4, and TEK. Although we highlighted the role of ECs in cell-to-cell communication instead of pre-ECs, the contribution of pre-ECs in the advancement of OA could not be neglected. We found that specific receptors on OBs, including EPH, NOTCH, BMPR, NRP, PDGFR, TGFβR, fibronectin, collagen, and SPP1 (OPN), responded to signals derived from ECs, thus further elucidating the interaction between the osteogenic subgroups and ECs. Furthermore, EnOBs are related to angiogenesis and promote OB proliferation. StOBs express higher levels of fibrin and collagen. Moreover, OPN, which is directly related to mineralisation, is expressed only by MinOBs.

In conclusion, our scRNA-seq analysis results provided a clearer and more consistent definition of the cellular components of human subchondral bone in OA. Specifically, two novel populations of ECs and three subpopulations of OBs, as well as the intercellular interaction network between these subpopulations, were identified. Our analysis provides new insights into the physiological and pathological behaviours of subchondral bone in OA pathogenesis, which may contribute to novel therapeutic strategies in the future.

## Materials and Methods

Human subchondral bone samples were collected during total knee arthroplasty operations. Bone samples were cut into 1–2 mm pieces and then digested in 0.2% type II collagenase (17101015, Thermo Fisher Scientific, MA, USA) for 2 h. Cells were then collected after red blood cell lysis.

### Raw data processing and quality control

The Cell Ranger software pipeline (version 3.1.0) provided by 10× Genomics was used to demultiplex cellular barcodes, map reads to the genome and transcriptome using the STAR aligner, and down-sample reads as required to generate normalized aggregate data across samples, producing a matrix of gene counts versus cells. We processed the unique molecular identifier (UMI) count matrix using the R package Seurat (version 3.0). To remove low quality cells and likely multiplet captures, which is a major concern in microdroplet-based experiments, we apply a criteria to filter out cells with UMI/gene numbers out of the limit of mean value +/- 2 fold of standard deviations assuming a Guassian distribution of each cells’ UMI/gene numbers. Following visual inspection of the distribution of cells by the fraction of mitochondrial genes expressed, we further discarded low-quality cells where a certain percentage of counts belonged to mitochondrial genes. Library size normalization was performed in Seurat on the filtered matrix to obtain the normalized count.

Top variable genes across single cells were identified using the method described in Macosko et al. Briefly, the average expression and dispersion were calculated for each gene, genes were subsequently placed into several bins based on expression. Principal component analysis (PCA) was performed to reduce the dimensionality on the log transformed gene-barcode matrices of top variable genes. Cells were clustered based on a graph-based clustering approach, and were visualized in 2-dimension using tSNE. Likelihood ratio test that simultaneously test for changes in mean expression and in the percentage of expressed cells was used to identify significantly differently expressed genes between clusters. Here, we use the R package SingleR, a novel computational method for unbiased cell type recognition of scRNA-seq to infer the cell of origin of each of the single cells independently and identify cell types.

Differentially expressed genes(DEGs) were identified using the Seurat package.p value < 0.05 and |log2foldchange| > 1 (or |log2foldchange| > 0.58) was set as the threshold for significantly differential expression. GO enrichment and KEGG pathway enrichment analysis of DEGs were respectively performed using R based on the hypergeometric distribution.

### Pseudotime analysis

Pseudotime analysis was performed with Monocle2 to determine the dramatic translational relationships among cell types and clusters. Further detection with the Monocle2 plot_pseudotime_heatmap function revealed the key role of a series of genes in the differentiation progress. Signifcantly changed genes were identified by the differential GeneTest function in Monocle2 with a q-value < 0.01.

### Cell–cell communication analysis with CellPhoneDB 2

CellPhoneDB 2 is a Python-based computational analysis tool developed by Roser Vento-Tormo et al, which enables analysis of cell–cell communication at the molecular level. A website version was also provided for analysis of a relatively small dataset (http://www.cellphonedb.org/). As described above, 606 single cells that were clustered into 5 cell types were investigated using the software to determine interaction networks. Interaction pairs including chemokines, ephrin receptor family, NOTCH family, cytokines and integrins and have p-values < 0.05 returned by CellPhoneDB, were selected for the evaluation of relationships between cell types.

### SCENIC analysis

SCENIC is a new computational method used in the construction of regulatory networks and in the identification of different cell states from scRNA-seq data. To measure the difference between cell clusters based on transcription factors or their target genes, SCENIC was performed on all single cells, and the preferentially expressed regulons were calculated by the Limma package. Only regulons significantly upregulated or downregulated in at least one cluster, with adj. p-value < 0.05, were involved in further analysis.

### Animal models

Destabilization of the medial meniscus (DMM) or sham operations were conducted bilaterally on 6-week-old male C57B6J mice (n=5 in each group). Samples were harvested at 0, 2, 4 and 8 weeks after surgery. Mice of 0 weeks were conducted with sham operation. Hearts were perfused with PBS and 4% PFA successively in order to fix the antigens. Bilateral knee joints were harvested and fixed in 4% PFA for 24 hours, then went through micro-CT scan (60kV, 50μA, 10 um pixel).

### Pathological staining

Samples were decalcified in 10% EDTA for two weeks, paraffin sections of 6 μm were then prepared for subsequent experiments: HE, safranin O and fast green (SOFG) and immunofluorescence (IF) staining. HE and SOFG staining experiments were conducted under manufacturer’s instructions (Beyotime, C0105M; Solarbio, G1371). Antibodies utilized during IF staining: Rat anti Mouse PECAM1 (Thermo fisher, 140311-82); Rabbit anti Mouse KDR (Abcam, ab11805); Rat anti Mouse OCN (Takara, M188); Rabbit anti Mouse MGST1 (Abcam, ab131059); Rabbit anti Mouse GFPT2 (Abcam, ab190966); Rabbit anti Mouse WIF1 (Santa, sc-373780); Goat anti Rat secondary antibody (Abcam, ba150165); Goat anti Rabbit secondary antibody (Abcam, ab150080). Nuclei were marked by DAPI (Beyotime, C1002).

### Statistical analysis

Statistical calculations were performed using R package and GraphPad Prism (version 9.3). The results of microCT and staining are presented using line charts. The two-way ANOVA was applied to identify differences between groups in statistical graphs. Results were considered statistically significant when *p* < 0.05.

## Acknowledgements

We thank OE biotech company (Shanghai, China) for the support of bioinformatics analysis. This work is supported by National Key R&D Program of China (2018YFC2001500), National Natural Science Foundation of China (82172098).

## Author contributions

Conceptualization: JCS, XC, KX; Methodology: YH, JC, HL, SCW; Investigation: YH, KX, HL; Visualization: KX, YH; Supervision: JCS, LHC; Writing—original draft: YH, KX; Writing—review & editing: JCS, XC.

## Ethics approval

Protocol approved by the Ethical Committee of Shanghai University (ECSHU 2021-146).

