## Supplemental for "Single-cell RNA-sequencing analysis reveals the molecular mechanism of subchondral bone cell heterogeneity in the development of osteoarthritis"

**This PDF file includes:**

Figs. S1 to S10
Tables S1 to S2

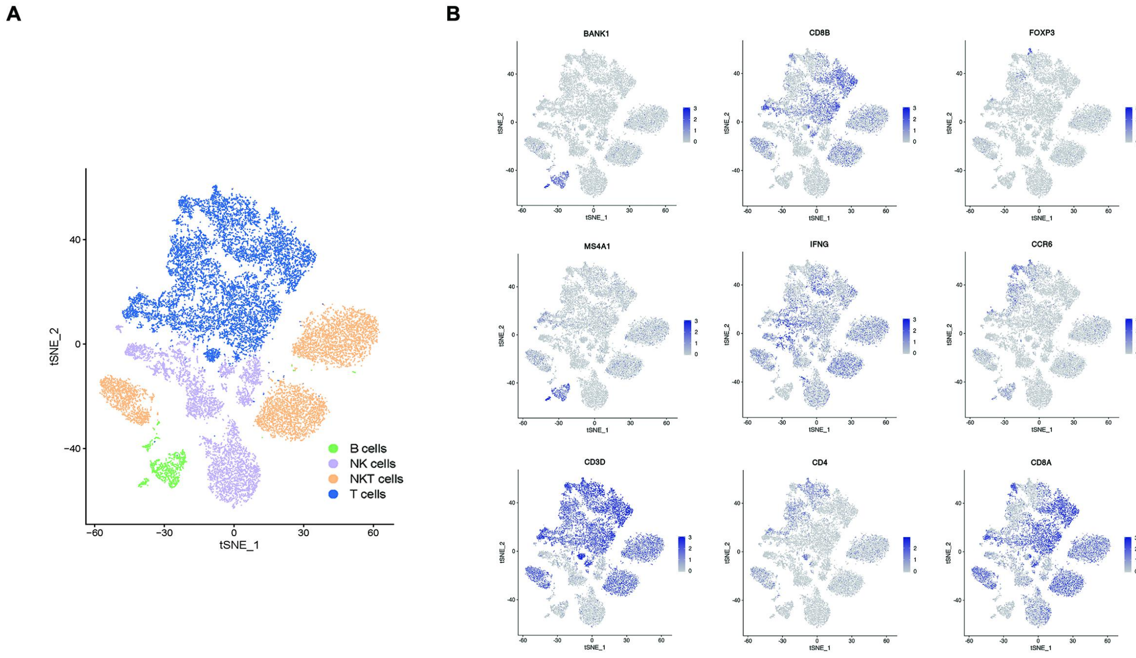


**Fig. S1. Single-cell profiling of human OA subchondral bone immune cells.** (**A**) t-SNE plots of Immune cells colored by cluster. (**B**) Dot plots showing the expression of indicated markers for each cell types on the t-SNE map.

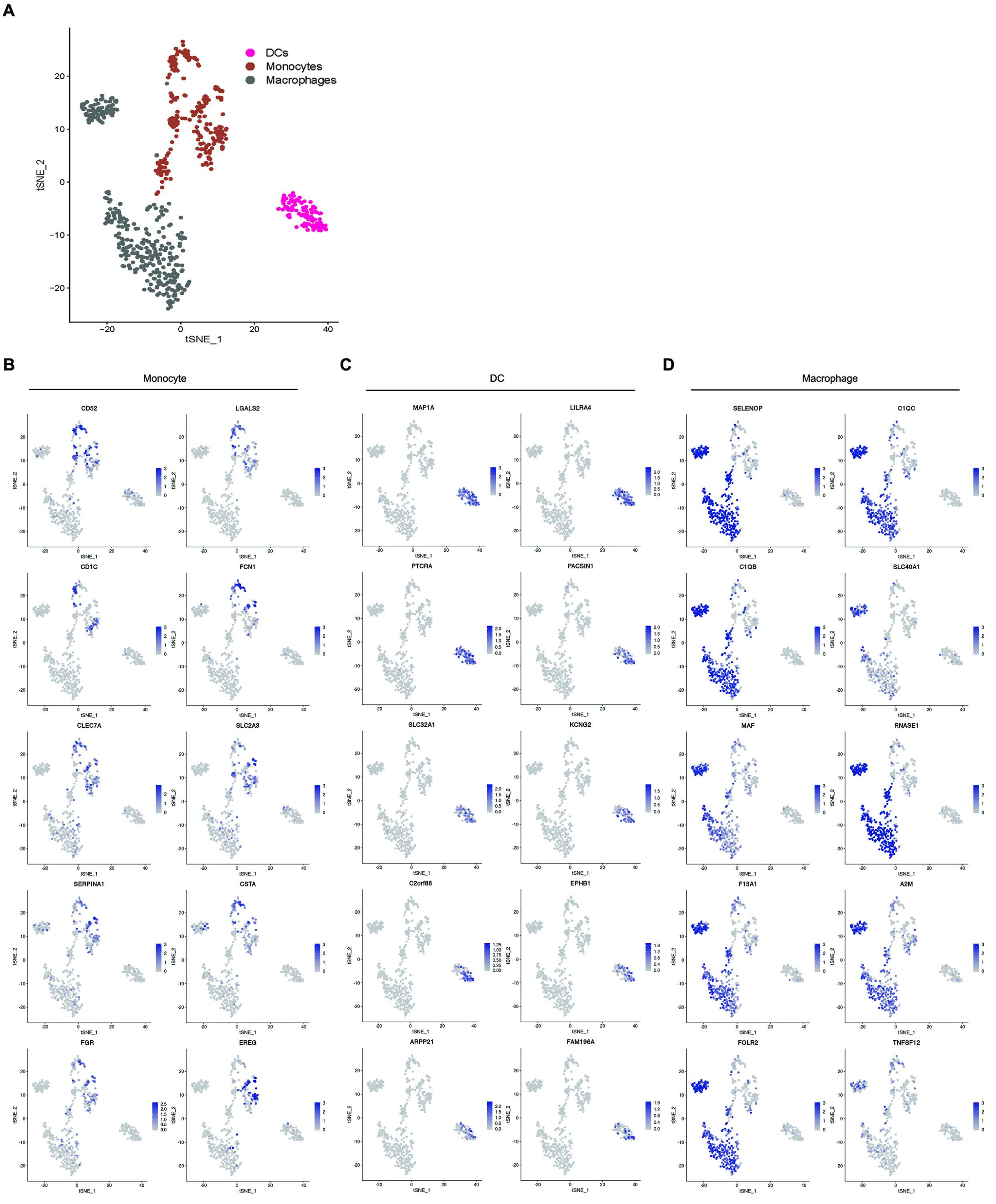


**Fig. S2. Single-cell profiling of human OA subchondral bone myeloid cells.** (**A**) t-SNE plots of myeloid cells colored by cluster. (**B**-**D**)Dot plots showing the expression of indicated markers for each cell types on the t-SNE map.

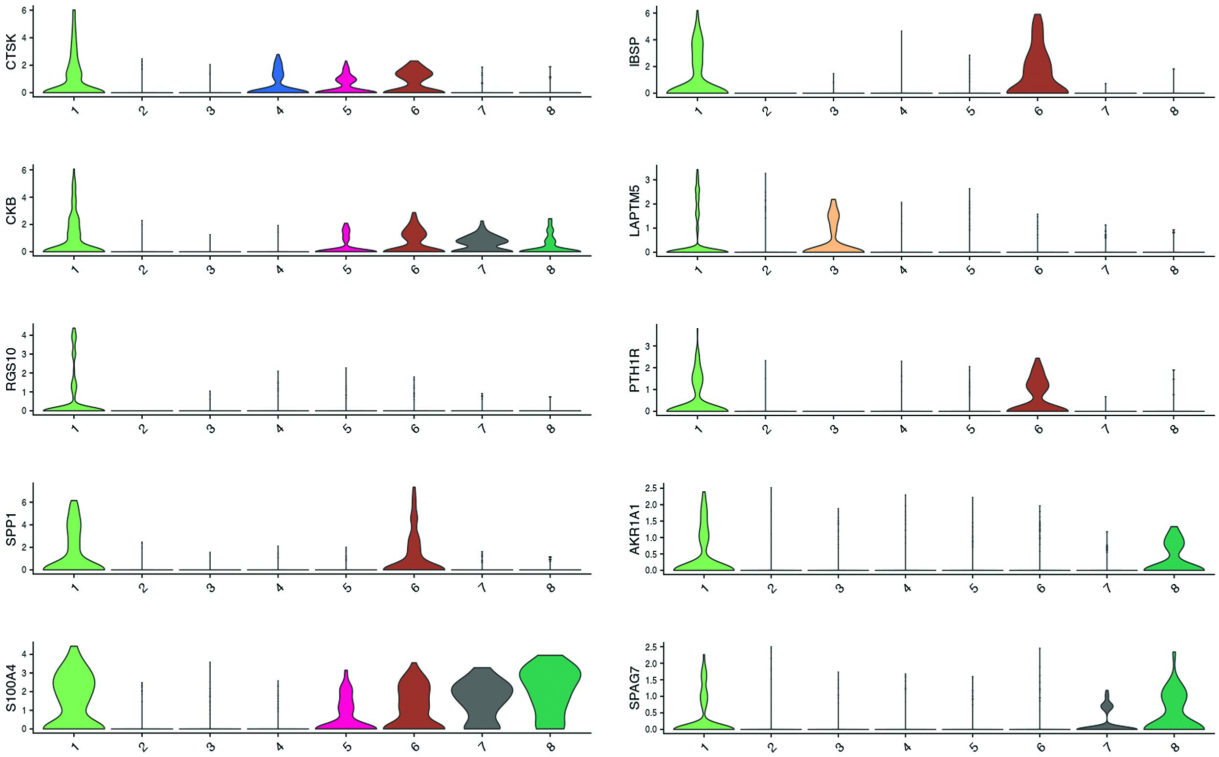


**Fig. S3. Violin plots showing expression levels of indicated markers for Cluster 1**

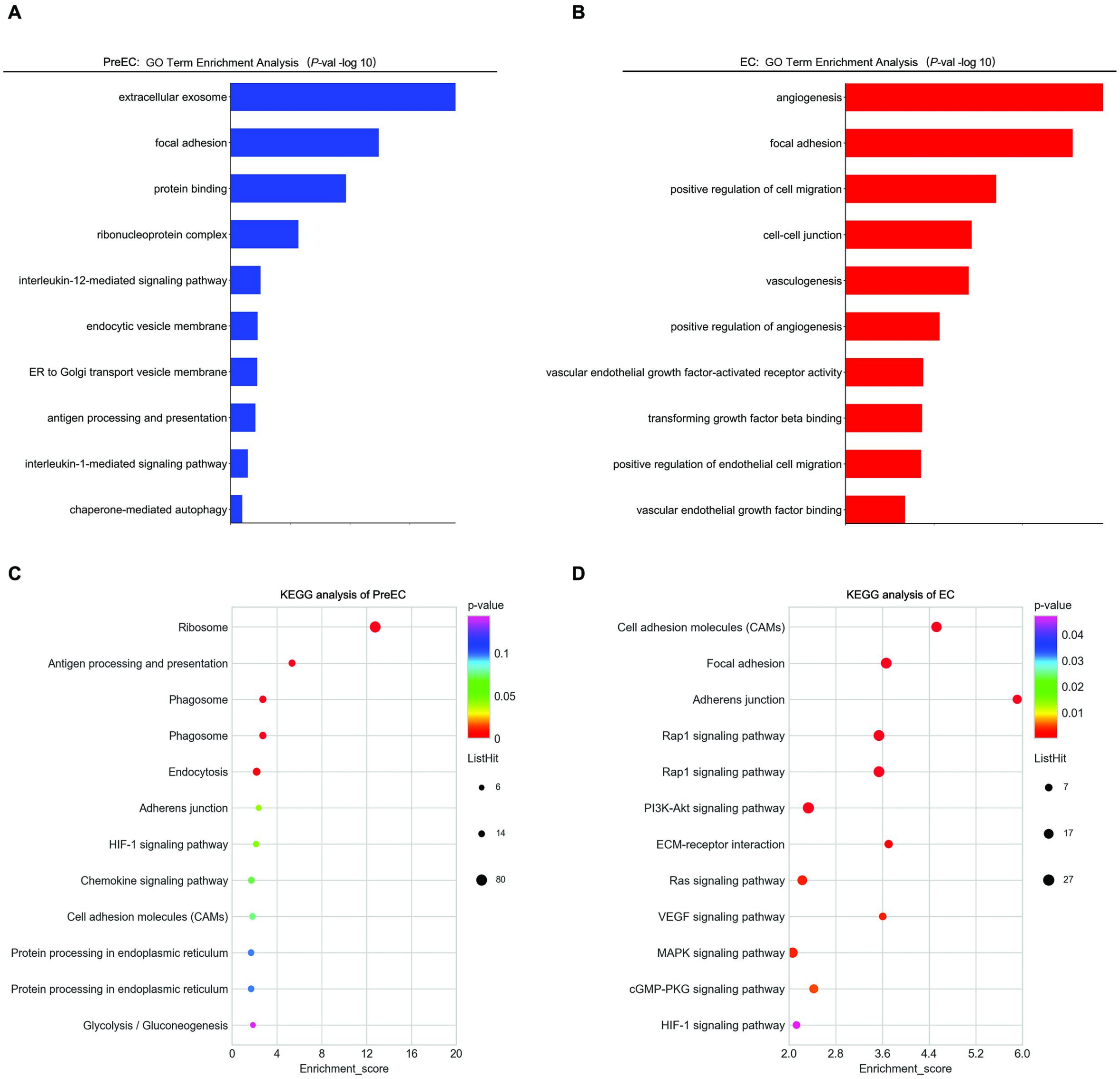


**Fig. S4. Identification of Pre-ECs and ECs.** (**A**)Enriched GO functions of upregulated genes in Pre-ECs. (**B**)Enriched GO functions of upregulated genes in ECs. (**C**)Enrichment plot from KEGG pathway analysis for Pre-ECs. (**D**)Enrichment plot from KEGG pathway analysis for ECs. Type or paste caption here. Create a page break and paste in the Figure above the caption.

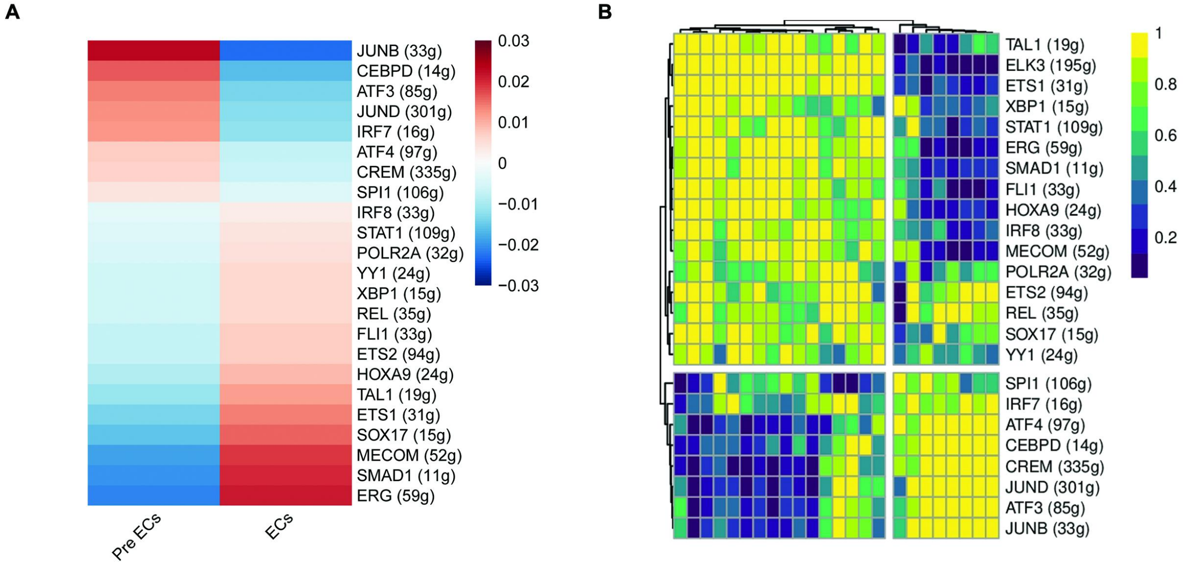


**Fig. S5.The difference between Pre-ECs and ECs in transcriptional regulation.** Single-cell regulatory network inference and clustering (SCENIC) analysis showing distinct regulons between ECs and Pre-ECs.The heatmap listing only the regulons with significant differences.

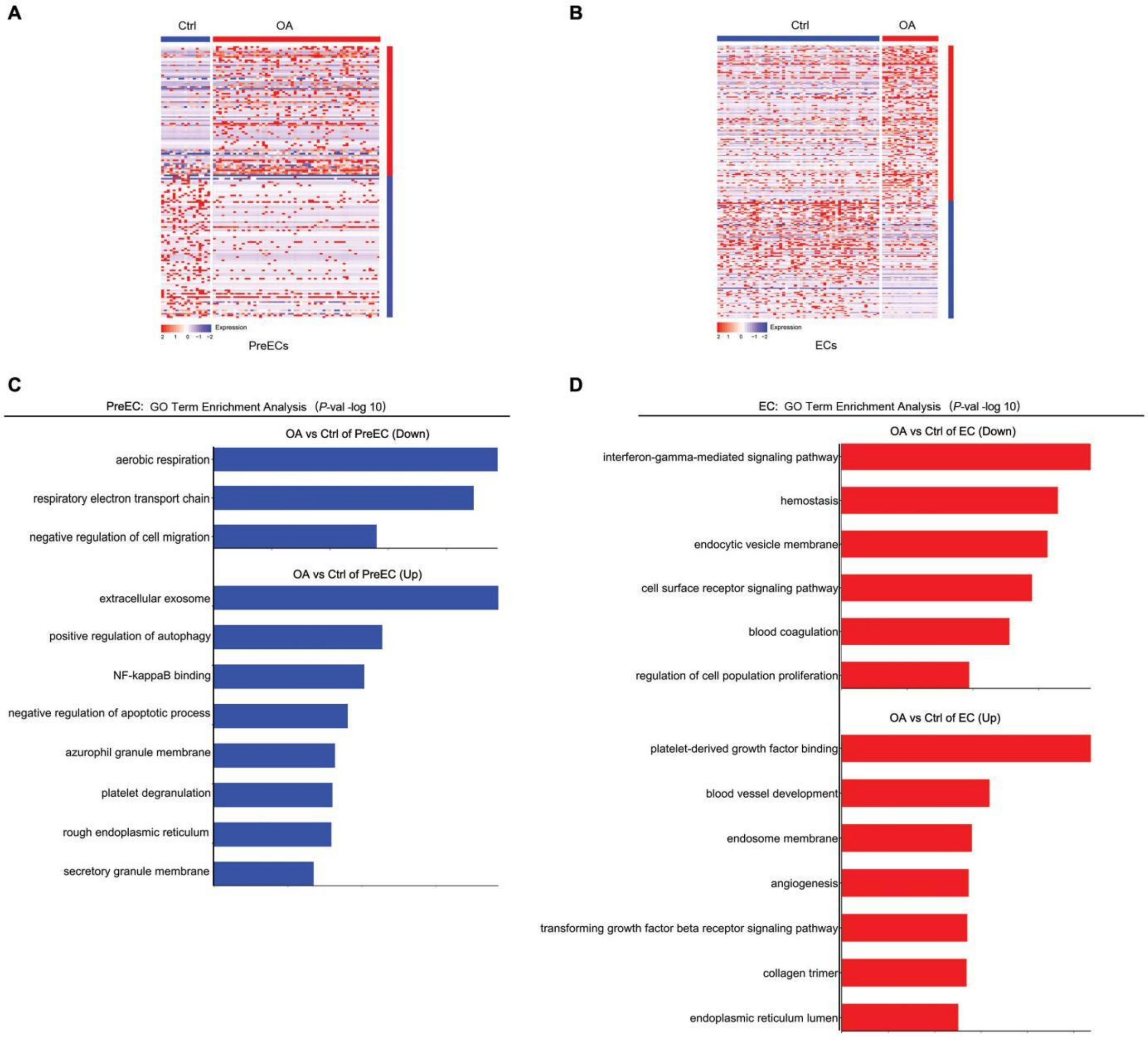


**Fig. S6. Identification of OA group and Ctrl group in endothelial cells.** (**A**)Heatmap of DEGs between OA group and Ctrl group in Pre-ECs and ECs. (**B**)GO functions enrichment analysis of OA versus ctrl upregulated genes in Pre-ECs and ECs

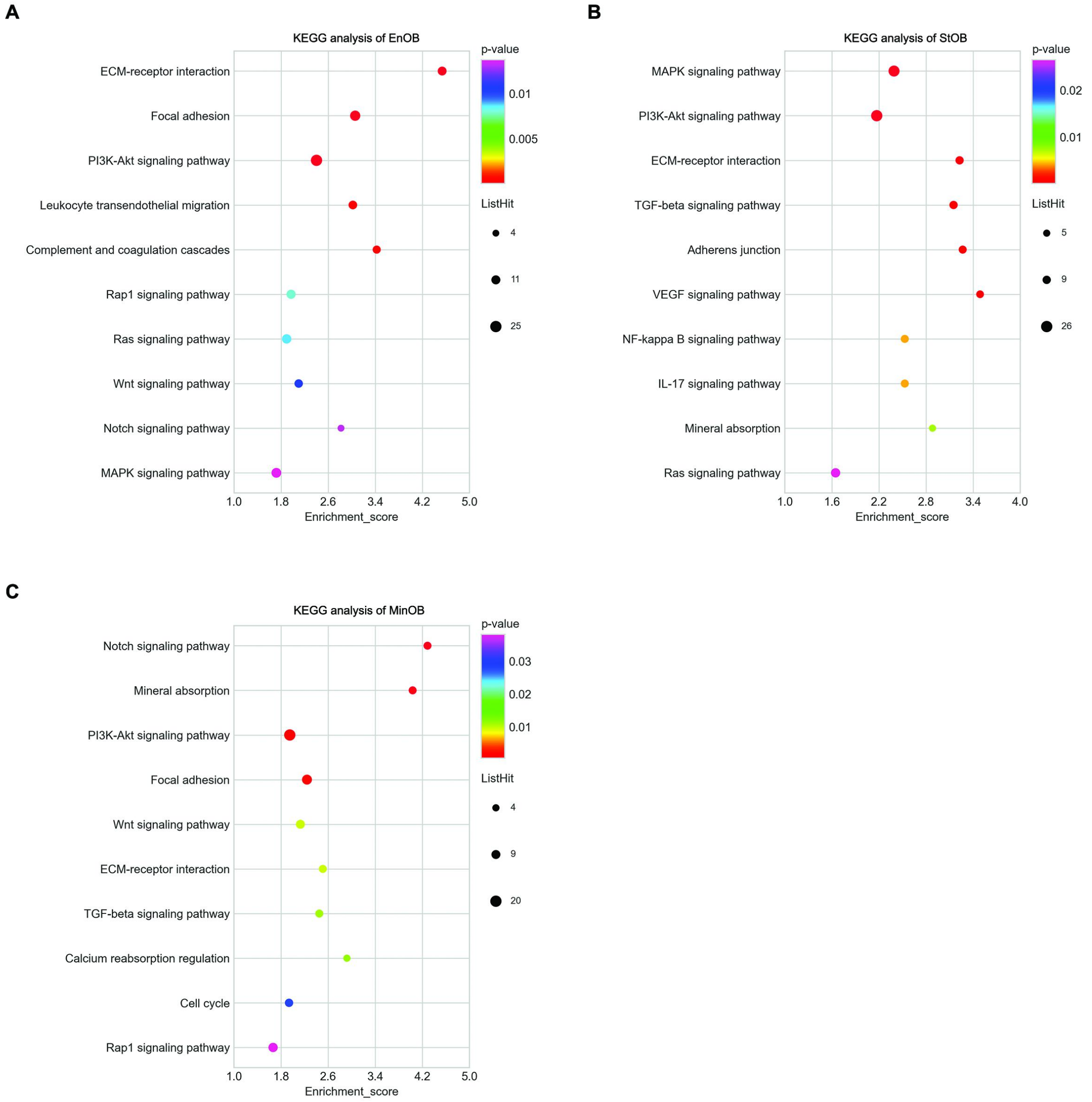


**Fig. S7. KEGG showing enrichment of pathways among EnOBs, StOBs and MinOBs.** (**A**)Enrichment plot from KEGG pathway analysis for EnOBs. (**B**)Enrichment plot from KEGG pathway analysis for StOBs. (**C**)Enrichment plot from KEGG pathway analysis for MinOBs

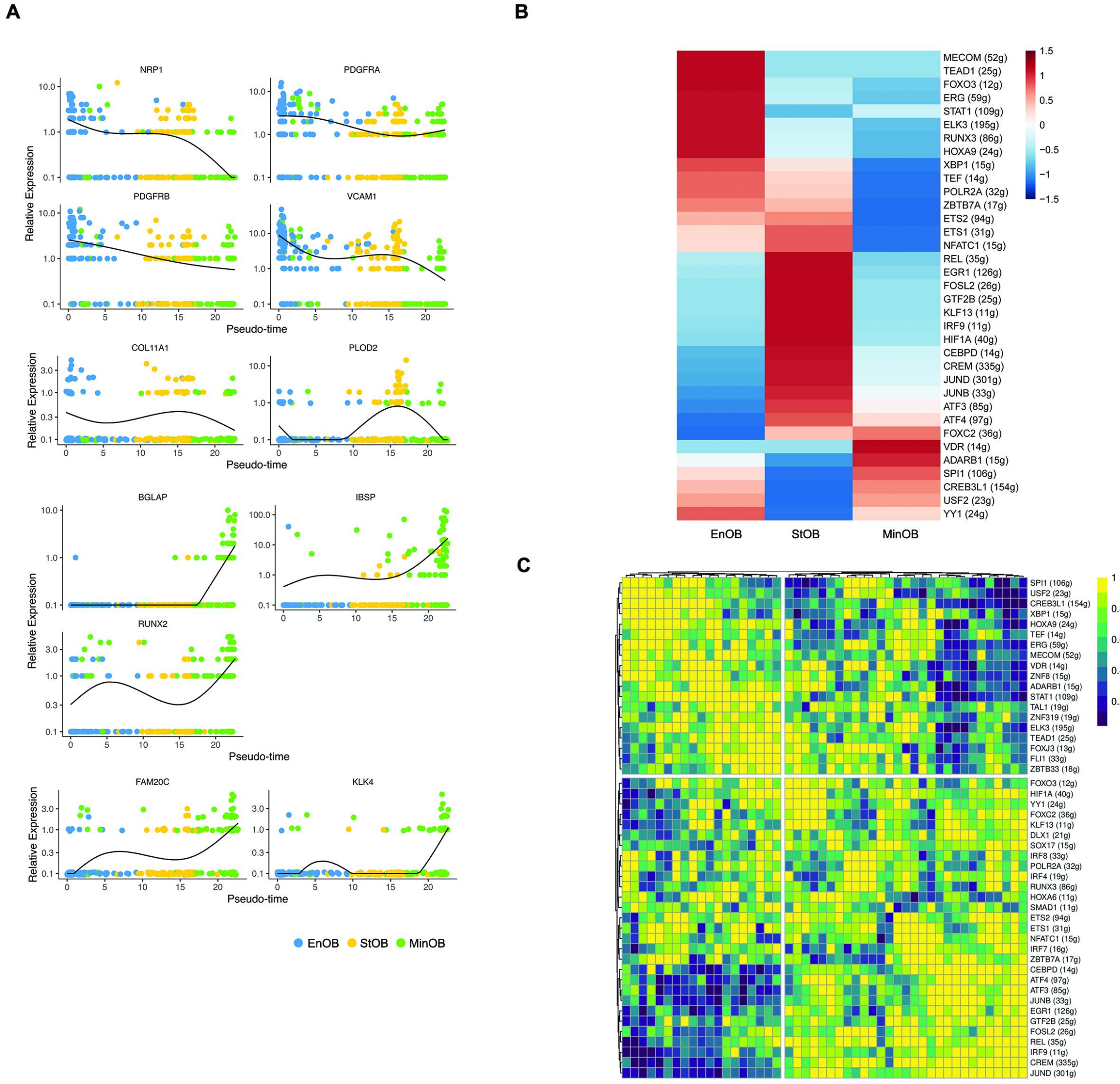


**Fig. S8. Identification of EnOB, StOB and MinOB.** (**A**)Pseudotemporal expression dynamics of marker gene in EnOBs,StOBs and MinOBs. All single cells in the EnOBs,StOBs and MinOBs cell lineage are ordered based on pseudotime. (**B-C**) Single-cell regulatory network inference and clustering (SCENIC) analysis showing distinct regulons among EnOBs,StOBs and MinOBs.The heatmap listing only the regulons with significant differences.

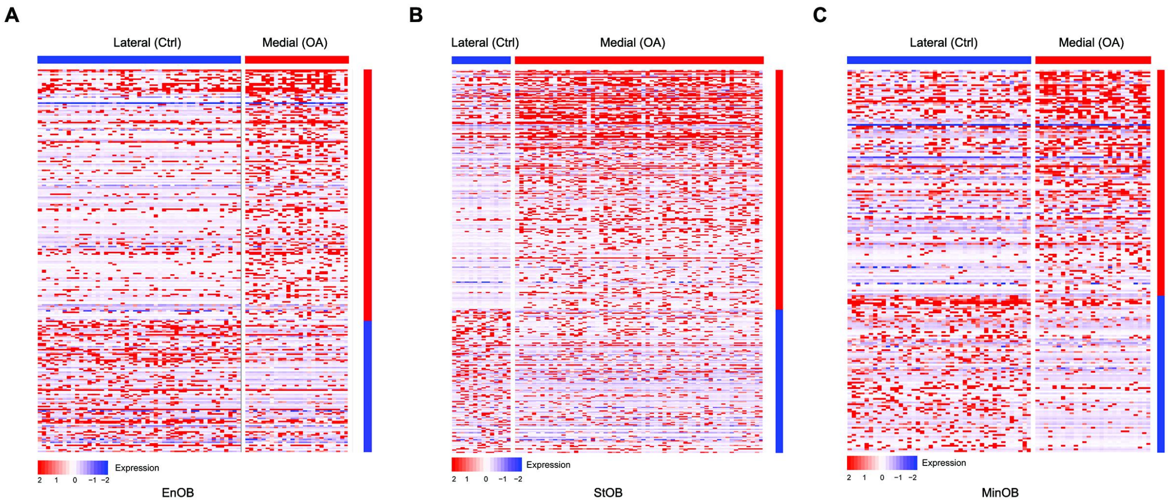


**Fig. S9. Identification of OA group and Ctrl group in osteoblasts.** (**A**)Heatmap of DEGs between OA group and Ctrl group in EnOBs. (**B**)Heatmap of DEGs between OA group and Ctrl group in StOBs. (**C**)Heatmap of DEGs between OA group and Ctrl group in MinOBs

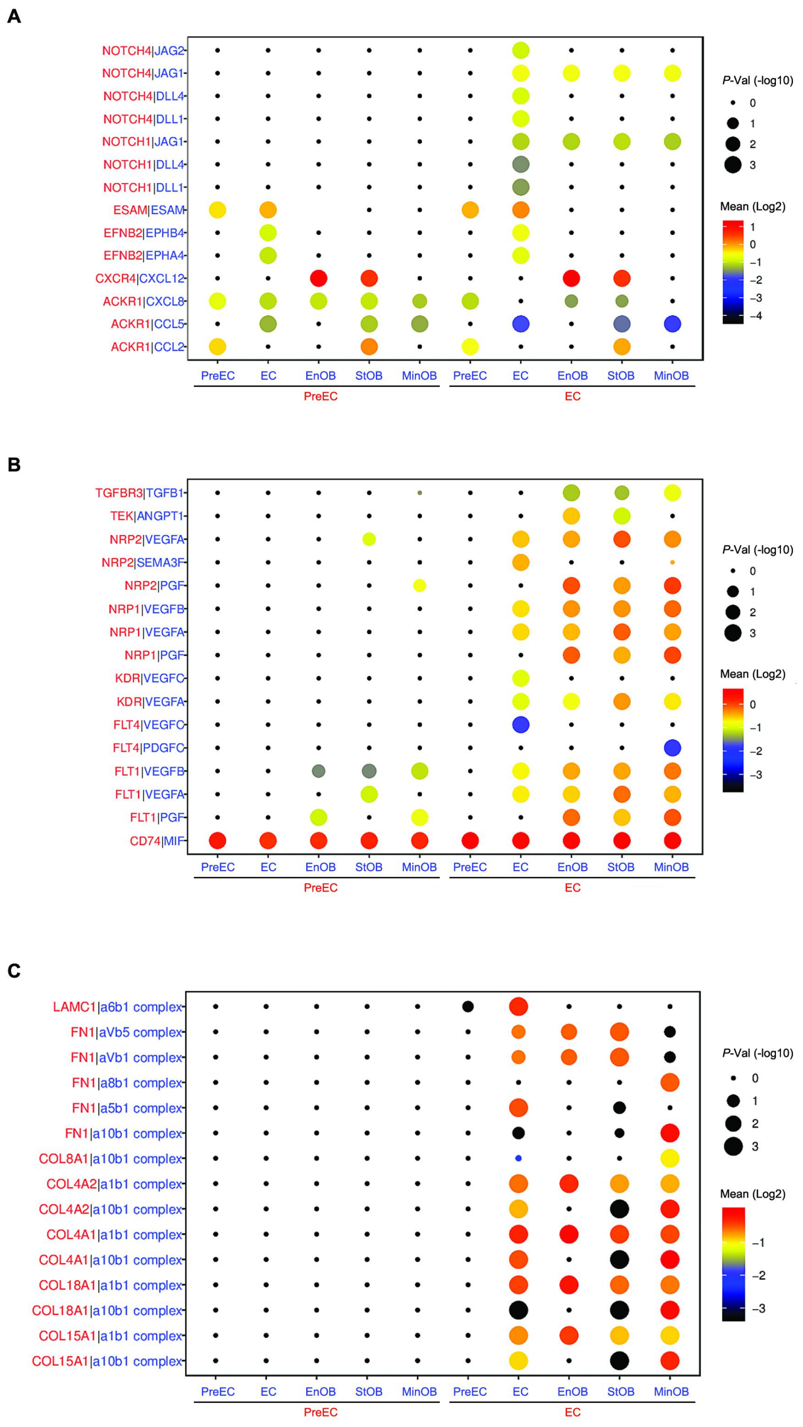


**Fig. S10. Vascular endothelial cell and osteoblast subtype interaction.** CellPhoneDB analysis showing the number of ligand-receptor interactions between endothelial cell subpopulation and osteoblast subpopulation. Bubble plots show ligand-receptor pairs of cytokines (**A**) , growth factors (**B**) and integrin (**C**) between endothelial cells subpopulation.

**Table S1. Clinical and demographic characteristics of the patients**

| | Patient 1 | Patient 2 |
|---|---|---|
| Gender | Female | Female |
| Age (years) | 74 | 72 |
| BMI | 26.89 | 29.14 |
| TKA | Right | Left |
| KSS score | 54 | 58 |
| Chronic conditions | Hypertension (grade Ⅱ) | Type 2 diabetes, hypertension (grade Ⅰ) |

**Table S2. Markers of each cell type**

| Cell type | Markers |
|---|---|
| B cell | CD79A, BANK1, MS4A1 |
| NK cell | GZMB, NKG7 |
| NKT cell | NKG7, CD3D |
| T cell | CD3D, CD3G |
| DC | LILRA4, PTCRA |
| Monocyte | CSTA, FCN1 |
| Macrophage | CD14, CD68, CSF1R, C1QC, F13A1 |
| EC | PECAM1, CLDN5 |
| MSC | MCAM |
| OB | RUNX2, CDH11 |
